# Mapping fitness landscapes to interpret sex allocation in hermaphrodites

**DOI:** 10.1101/2024.07.23.604733

**Authors:** Kai-Hsiu Chen, John R. Pannell

**Author notes:** X (formerly Twitter): @kaihsiu_chen (Kai-Hsiu Chen).

## Abstract

Sex allocation theory predicts sex ratios of dioecious organisms, but it has been poor at explaining sex allocation in hermaphrodites in which the assumed trade-off between male and female functions is often obscure. Here, we apply sex-allocation theory to hermaphrodites by mapping components of seasonal reproductive success onto a fitness landscape defined by potentially independent measures of allocation to male and female functions on orthogonal axes. We find that peaks of reproductive success in a perennial hermaphroditic plant reflect the interactive effect of both male and female allocations on self-fertilization and the effects of inbreeding depression. The rugged landscape corresponds well to the complex pattern of sex allocation observed in natural populations in which individuals produce a mix of male and bisexual flowers and express a type of gender diphasy. Our approach may help to interpret common complexities of sex allocation in hermaphroditic plants and animals.

## Introduction

The theory of sex allocation has provided a simple and powerful framework for understanding the reproductive strategies of species with two different gamete sizes^1^. The theory explains when a species should be dioecious versus hermaphroditic^2^, what sex ratio dioecious species should have under different scenarios of inbreeding, kin competition or conflict^3,4^, whether sex should be determined by genes or by the environment^5^, what proportion of reproductive resources hermaphrodites should allocate to their two sexual functions^2,6^, whether hermaphrodites should allocate to both sexes simultaneously or sequentially^7^, and whether (and how) their allocation strategy should depend on their size^8,9^. But whereas sex-allocation theory has been relatively successful in predicting the sex ratio and timing of allocation to sons versus daughters in species with two separate sexes^10^, it has been notoriously difficult to test in populations that include hermaphrodites^11–16^ (but see sex change in hermaphroditic animals^17^).

Applying sex-allocation theory to hermaphrodites has been unsuccessful for several reasons. First, the theory is based fundamentally on the assumption that individuals must divide a given reproductive resource between their male and female functions in a zero-sum game, with a simple allocation trade-off between the two sexes^1,2,6^. This assumption is straightforward in dioecious species in which the sex allocation is simply the progeny sex ratio^4,18^. However, empirical studies on hermaphroditic species have found it difficult to identify the resource that hermaphrodites must divide between their male and female functions – and even to define what ultimately constitutes allocation to each function, e.g., whether it should include costly ancillary structures that promote mating success^12,14,15^. Size-dependent sex-allocation theory recognizes a likely important dimension of individual variation in resource availability, but it does not tell us which resource to measure or precisely how the resource should vary with an individual’s size^8^. Ignorance of the relation between size and resource availability may be one of the reasons for our failure, with few exceptions, to uncover male-female allocation trade-offs – even when size is statistically accounted for^19,20^. Moreover, sex-allocation trade-offs may also be obscured by trade-offs between reproduction and other allocation sinks such as growth or defense^21–24^.

Second, testing sex-allocation theory for hermaphrodites requires an accurate estimate of the fitness gained through each sexual function, but we have few estimates of both male and female reproductive success in natural populations, especially in plants^25–27^. Estimates of the female reproductive success of hermaphrodites have been easier than those of male reproductive success, but they have usually been made in terms of the number of seeds or fertilized eggs without accounting for establishment probability or viability^12^. In self-compatible hermaphrodites, self-fertilization is known to strongly affect progeny establishment through inbreeding depression^28^. Yet although some sex-allocation models incorporate the effects of selfing and inbreeding depression^2,29^, to our knowledge there are still no empirical tests of these models or evaluations of how fitness gains are affected by inbreeding.

Third, tests of sex-allocation theory have been largely limited to considering allocation at the individual level^30–32^, yet in modular organisms such as plants, sex allocation often varies in complex ways among their modules^33,34^. While most flowering plants produce bisexual flowers with both male and female sexual organs, the individuals of many species vary in their sex allocation among flowers or reproductive modules. Such heteromorphic strategies include ‘monoecy’ (the simultaneous production of separate male and female flowers), and ‘andromonoecy’ (the production of bisexual and male flowers)^35^. In many functionally hermaphroditic species, individuals also vary their sex allocation over time, either within a season in terms of phenology (e.g., flowers may be protandrous or protogynous), or among seasons (e.g., species showing ‘gender diphasy’, in which individuals shift from being male to female or hermaphrodite over the course of their lives)^36,37^. Some models of sex allocation seek to predict the optimal allocation strategy for ‘temporal’ modules of an organism, i.e., how it should allocate its reproductive resources at different times or ages, often corresponding to different sizes^8,21,24,38^, but we know of only one attempt to use sex allocation theory to understand the evolution of separate male and female modules on the same individuals^39^, and no studies have attempted to test such ideas empirically.

Here, we overcome some of the limitations of applying sex-allocation theory to hermaphrodites by using a different approach that sidesteps the trade-off assumption (whether it is upheld or not). Our approach involves mapping male and female fitness components on orthogonal axes of a two-dimensional fitness landscape. Although mapping reproductive success to genotypes with continuous traits in a fitness landscape has been common in evolutionary biology^40–42^, to our knowledge this approach has not previously been applied in empirical studies of sex allocation in hermaphrodites. We illustrate our approach by using it to interpret the complex hermaphroditic allocation strategy of the self-compatible perennial herb, *Pulsatilla alpina* (L.) Delarbre (Ranunculaceae), which displays both andromonoecy and a type of gender diphasy^43,44^. This variation demands a functional explanation, but it also presents an opportunity to consider the fitness implications of a wide range of allocation phenotypes at the flower level. While sex allocation is already highly variable among *P. alpina* flowers^45^, we further enhanced this variation by removing stamens from a subset of flowers (Figure 1). This manipulation allowed us to infer the fitness implications of producing female flowers, which do not occur naturally in the species (and to offer an explanation for why). By focusing on a population of *P. alpina* comprising mostly single-flowered individuals, we could use genetic markers to attribute paternity to individual flowers and their phenotypes, which is not possible for multi-flowered individuals.

**Figure 1.**
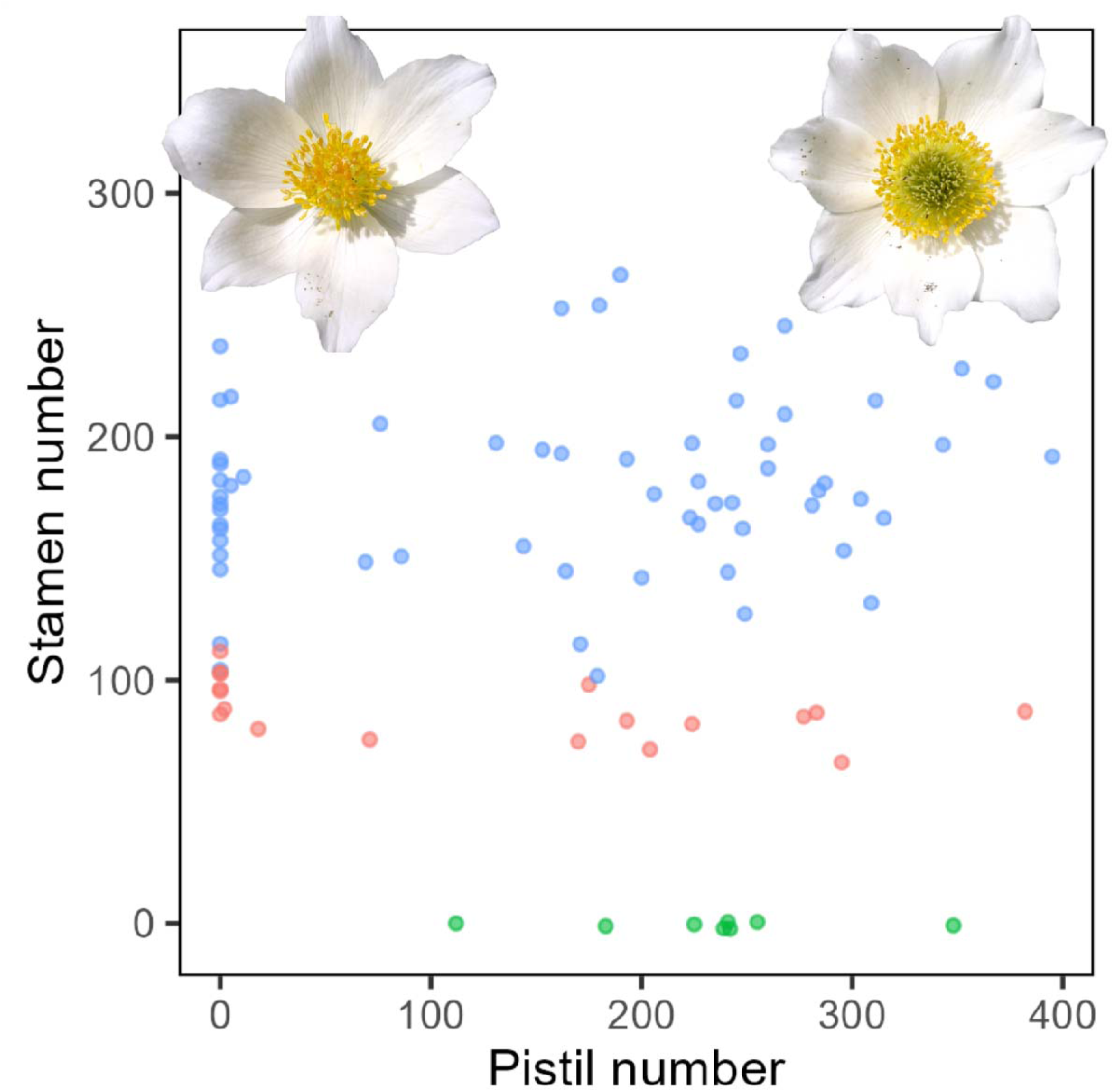
Morphological space of stamen and pistil number of the single-flowered individuals after stamen removal treatments. Green, orange, and blue points represent all-stamen-removed, half-stamen-removed, and intact flowers, respectively (*N* = 8, 18, and 61 respectively). Data points were jittered along the y-axis.

Although fitness is ultimately a property of individuals, by assessing the contribution to fitness through seed production and siring success at the modular, floral, level, we could begin to ask why a plant might produce both male and bisexual flowers with strongly different sex allocation. How, for instance, does the production of a male flower contribute to an individual’s marginal fitness gain compared to the production of a hermaphroditic (or female) flower in its place? Such an evaluation is presumably the general basis of allocation decisions taken by plants with heteromorphisms such as andromonoecy or monoecy, with each new module receiving resources for its male versus female functions depending on the context of, and prospective fitness contribution made by, that module and other modules of the same individial^46^ (we comment on limitations of using individuals with only single flowers in the Discussion).

We showcase our approach by modelling fitness components for flowers of *P. alpina* at the flower scale in terms of male and female allocations under two scenarios of inbreeding depression, both on their own as single variables and jointly. We first modelled estimates of male and female fitness in terms of components of sex allocation to characterize ‘fitness gain curves’ that relate fitness through each sexual function to allocation to that same function using univariate analysis. We then used multivariate selection gradient analysis to test how female and male allocations jointly affect fitness components. Finally, we map fitness components onto a two-dimensional fitness landscape. This mapping reveals fitness peaks that correspond closely to the complex sex-allocation strategy adopted by *P. alpina* in the wild, reflecting the interactive effects of male and female allocation beyond any simple trade-off. Our analyses provide new insights into the evolution of the complex reproductive strategies of a perennial and modular organism that may reinvigorate the empirical application of a conceptually powerful theory to hermaphrodites.

## Results

### Variation in floral sex allocation and components of reproductive success

We measured the male and female components of sex allocation for a total of 175 flowers, with 104 individuals producing a single flower and 31 individuals producing more than one flower (Figure S1). Prior to the stamen removal manipulations, 46 and 129 flowers were male and bisexual, respectively. Note that experimental stamen removal did not alter the flowering duration of the flowers^45^. We then estimated female, male, and total reproductive success for all individuals based on a paternity analysis using seeds sampled from all of the bisexual flowers in the population (*N* = 104 seed families; 25 bisexual flowers were aborted or missing). Each seed family produced 90.5 ± 55.4 mature seeds (mean ± SD; *N* = 104). We could assign paternity for 854 seeds to a single most likely father under a relaxed confidence interval (80%), corresponding to a 96% successful assignment rate for what amounted to around 9% of all mature seeds produced in the population. Our inferences are based on a sampling intensity greater than that of most recent studies of the mating system^47^ and phenotypic selection in hermaphroditic plants ^26,27,32,48–51^. Using selfing-rate estimates for seed families^45^ and adopting a value of inbreeding depression of δ = 0.95, which we estimated on the basis of comparisons of inbreeding coefficients between adults and seed progeny^52^, we inferred the mean female, male, and total reproductive success of single-flowered individuals to be 41.4 ± 48.6, 33.6 ± 41.5, and 75 ± 61.3, respectively (*N* = 87; flowers of 17 individuals were aborted or missing; Figure S2). These values compare with corresponding estimates of 65.8 ± 58.4, 57.9 ± 51.9, and 123.7 ± 88.3, respectively, under the assumption of a hypothetical scenario of δ = 0 (no difference in fitness between selfed and outcrossed progeny). Comparing fitness estimates between contrasting scenarios of inbreeding depression reveals that inbreeding depression may strongly affect inferences of selection on sex allocation and likely other traits.

### Female and male floral fitness gain curves

We estimated the shape of the male and female gain curves with a power function for individual flowers on the basis of reproductive success estimates for the 87 parents in our sample that produced a single flower (Figure 2). Regression of our estimate of female fitness on pistil production within flowers points to an accelerating female fitness gain curve (exponent *b* = 1.91 ± 0.3; mean ± se; significantly > 1.0; *P* < 0.01; Figure 2A and Table S1), confirming the result reported by Chen and Pannell^44^. Whereas under the hypothetical scenario that assumes no inbreeding depression, we infer a slightly saturating gain curve for the female function (*b* = 0.85 ± 0.16; Figure 2C), though the curve does not differ statistically from linearity (*b* is not significantly different from 1.0; *P* > 0.05; Table S1). In contrast to the inferred accelerating female fitness gain curve, our data point to saturating gain curves for male function, whether incorporating our estimate of inbreeding depression (*b* = 0.47 ± 0.38; Figure 2B) or not (*b* = 0.52 ± 0.27; Figure 2D) – though again neither of these two gain curve estimates differed significantly from linearity (*P* > 0.05; Table S1). Note that the high variance around the fitted male gain curves likely reflects the potential effects of cross-sex interactions not captured by univariate analysis or due to greater uncertainty around the estimates of male fitness.

**Figure 2.**
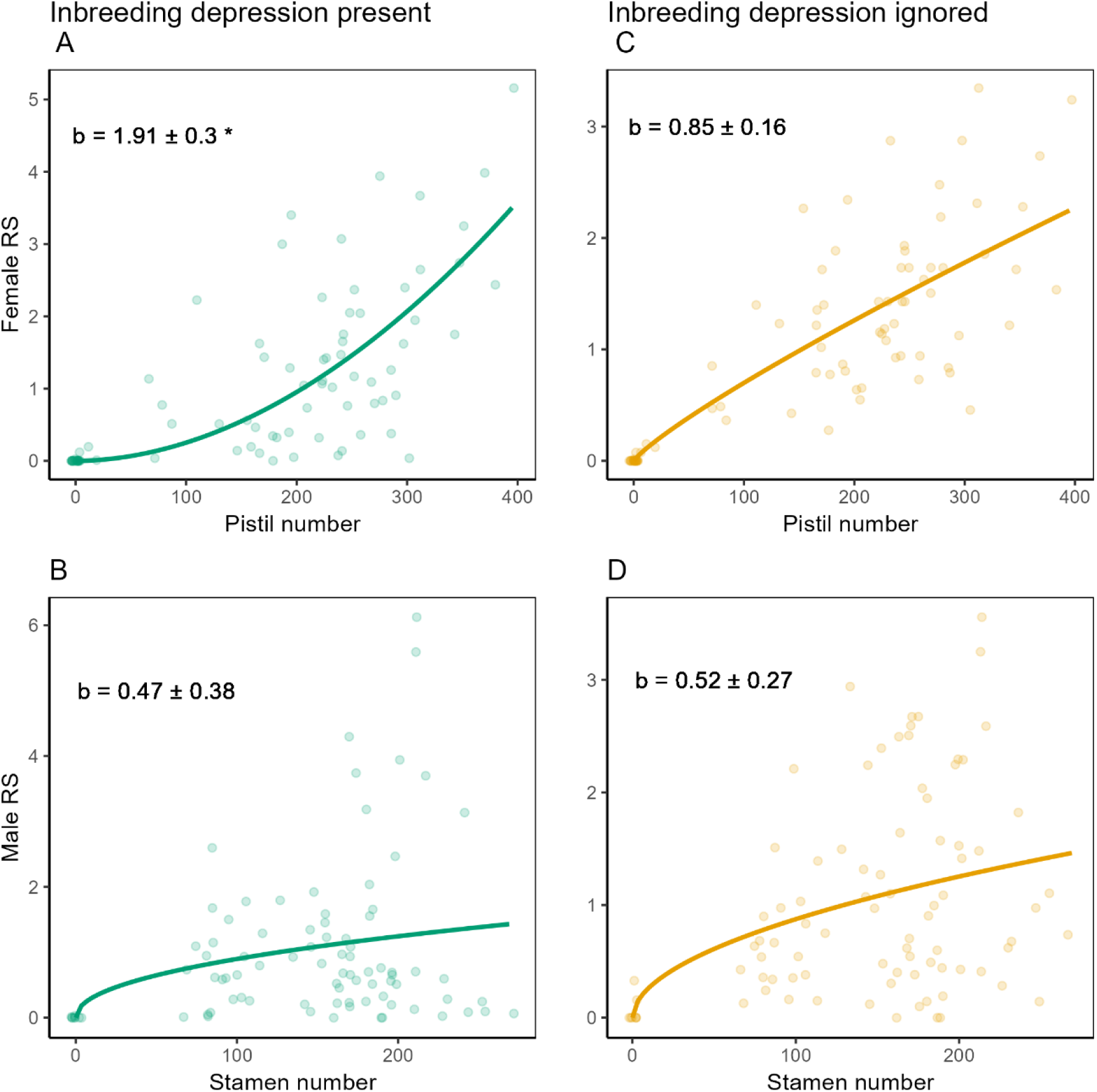
Fitness gain curves of female (upper panels) and male functions (bottom panels) at the floral level under the condition when inbreeding depression is taken into account (left-hand panels) or not (right-hand panels). Each point represents one individual with one flower (*N* = 87). The points were jittered to avoid overlapping. The shape of the gain curves was estimated by fitting exponential curves (see materials and methods for details) and the exponent *b* is shown in the figure with the standard error. An asterisk denotes that the curve was significantly non-linear.

### Linear, quadratic, and correlational selection gradients on female and male allocation

We further performed a multivariate selection gradient analysis^53^ on the single-flowered individuals (*N* = 87), which fully confirmed the above results: female reproductive success was a clearly accelerating function of allocation to pistils when incorporating inbreeding depression in our fitness estimates, but linear when inbreeding depression was ignored (Figure 3A). Similarly, male reproductive success was a mostly saturating function of allocation, though saturation was only significant for the hypothetical scenario of no inbreeding depression (Figure 3E and Supplementary Information Table S2). Our selection-gradient analysis for female reproductive success also points to mostly negative directional selection on male allocation when fitness estimates incorporate inbreeding depression (Figure 3D and Supplementary Information Table S2), as a result of ovule discounting^54^, because flowers with more stamens had a higher selfing rate^44^. For male reproductive success, our results point to stabilizing selection on female allocation, but only when inbreeding depression is ignored (Figure 3B and Supplementary Information Table S2). Total reproductive success depended only on female allocation when inbreeding depression was accounted for, but on both female and male allocation when ignoring it (Figure 3C and F; and Table S2). We could detect no correlational selection on female and male allocations (correlational selection gradients did not differ significantly from zero; Table S2).

**Figure 3.**
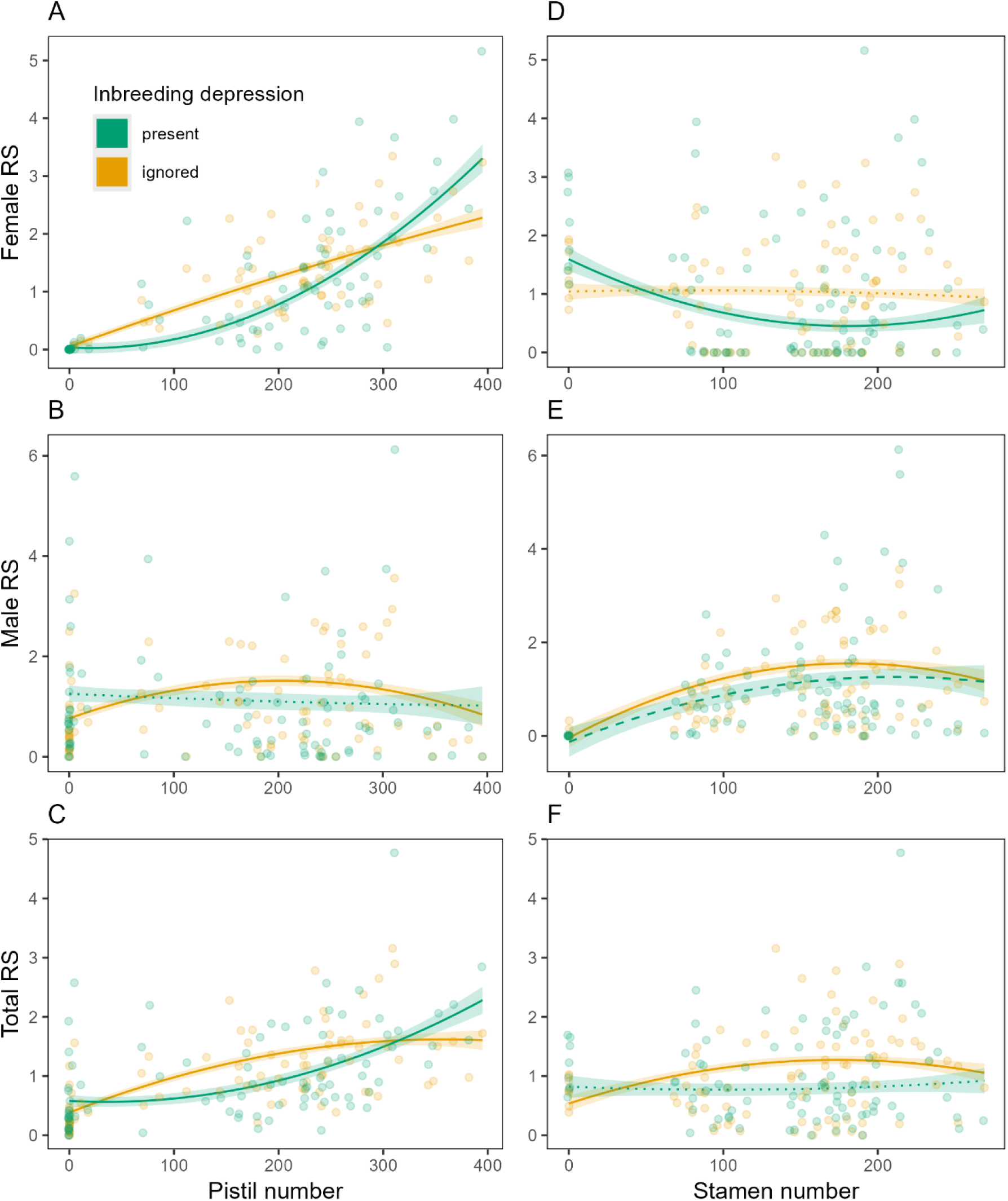
The dependence of female (upper panels), male (middle panels), and total (bottom panels) reproductive success (RS) on female and male allocation under the condition when inbreeding depression is taken into account (green lines) or ignored (yellow lines), estimated by selection gradient analyses (*N* = 87, see materials and methods for details; Table S2). The shaded ribbon indicates the range of one standard error of the partial regression curves. Regression lines of non-significant and marginally non-significant dependency of reproductive success on the sex function are shown in dotted and dashed lines, respectively. Raw data points were jittered to avoid overlapping. Panel **A** and **D** fully confirmed the results of a previous study based on the reproductive success of bisexual flowers only ^44^.

### Two-dimensional fitness landscapes for female, male, and total reproductive success

Maps of female, male, and total reproductive success on a two-dimensional sex-allocation landscape are presented in Figures 4A, B, and C, respectively. The two fitness peaks in the figure for total reproductive success correspond qualitatively closely to phenotypes of unisexual male and bisexual flowers, respectively (Figure 4C and S5). Standardized linear, quadratic, and correlational selection gradients, estimated using the *GSG* package for *R*^55^, are presented in Table S3. Note that we detected evidence for negative correlational selection gradients for total reproductive success when incorporating inbreeding depression in our fitness estimates (Table S3). Variance explained by fitted models for female, male, and total reproductive success were 79.9%, 58.2%, and 53.4%, respectively, when considering the influence of inbreeding depression, and 85.7%, 63.3%, and 67.5% when inbreeding depression was ignored.

**Figure 4.**
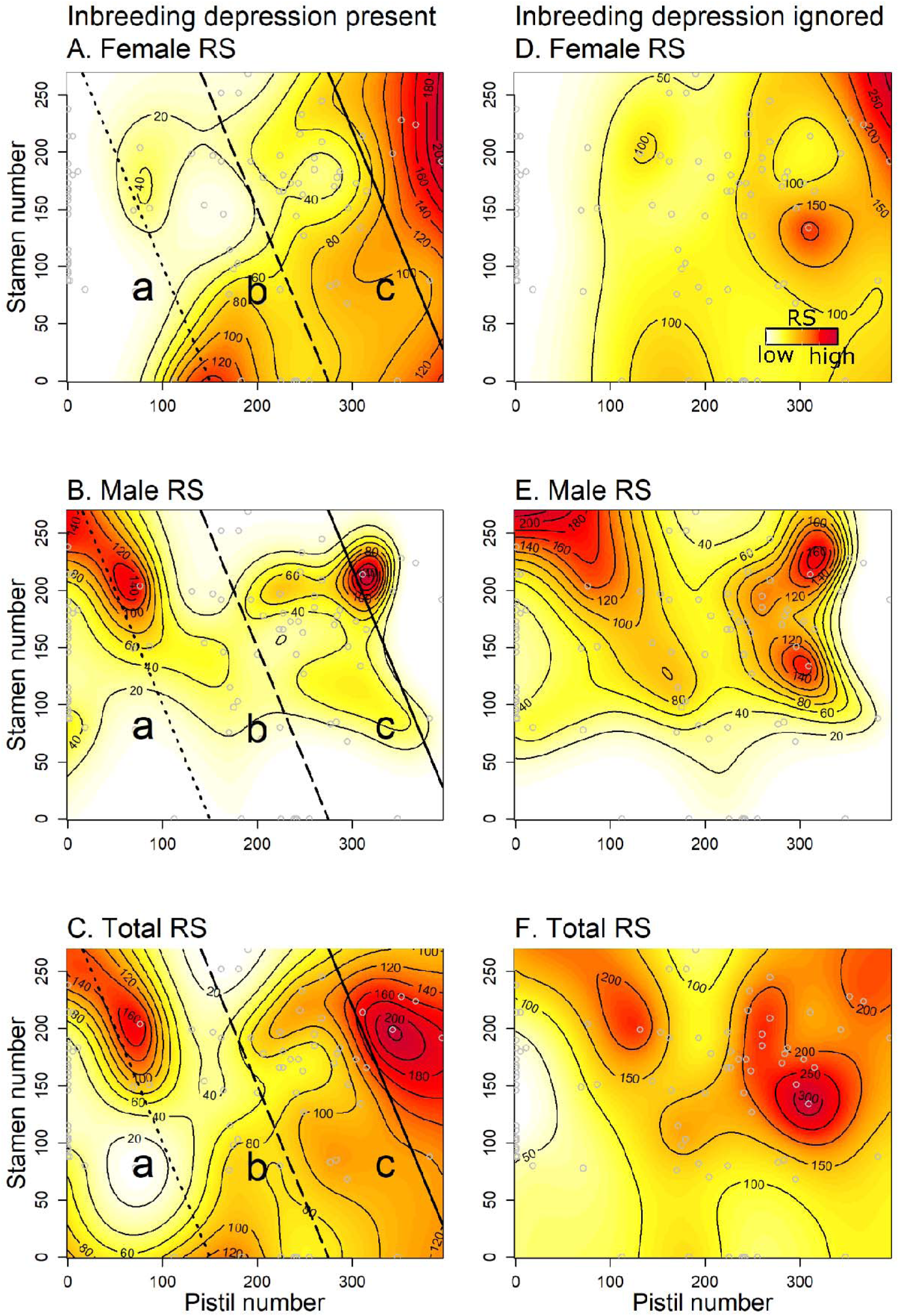
Representations of the fitness landscape for female, male, and total reproductive success (RS) as a function of pistil and stamen number in a flower under the condition when inbreeding depression is taken into account (left-hand panels) or ignored (right-hand panels), predicted by generalized additive models (*gam*) using 87 individuals with a single flower (grey circles). The colour gradient from white to red represents low to high predicted reproductive success in terms of the number of seeds. Hypothetical linear trade-off lines between male and female functions were depicted by dotted (line a), dashed (line b), and solid lines (line c) for individuals of low, medium, and high resource status, respectively (Panels **A**-**C**). Note that the slope of the trade-off lines is conceptual, because we do not know the linearity and the actual trade-off ratio relating female to male units of allocation. Individuals with a given amount of resource are only able to explore the left and bottom part of the trade-off line on the fitness landscape.

### Interpretation on gender-diphasy from the fitness landscapes

To aid interpretation of gender-diphasy in *P. alpina,* we drew hypothetical lines on the predicted fitness landscapes for strategies reflecting individuals of low, medium, and high resource status (when δ = 0.95); here, we assumed a linear sex-allocation trade-off with an arbitrarily chosen negative slope (Figure 4A, B, and C). We then plotted male, female and total fitness gain curves along each of these three hypothetical trade-off relations (following classic sex-allocation theory), with relative sex allocation computed as stamen number over the number of stamens and pistils together (Figures 5). Adopting either a unisexual male or a bisexual strategy yielded the highest total fitness returns for individuals with low (dotted line) and high (solid line) resource status, respectively (Figure 5C).

**Figure 5.**
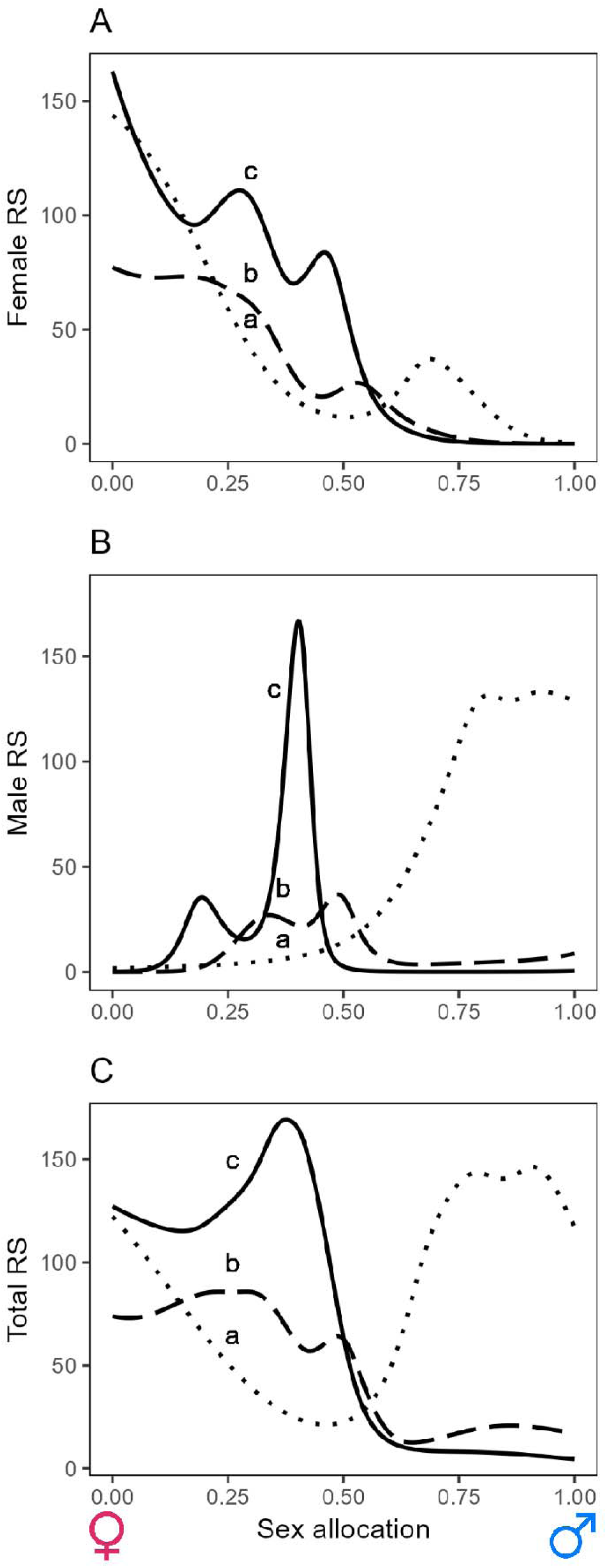
Conceptual figures demonstrating how female (**A**), male (**B**), and total (**C**) fitness gain curves depend on the resource status of an individual based on the study of *P. alpina*. Relationship of the projected reproductive success and sex allocation to the male function along the hypothetical trade-off lines a, b, and c for individuals of low, medium, and high resource status, respectively, were extracted from Figure 4 **A** - **C**. Sex allocation in terms of maleness was calculated by dividing the stamen number by the sum of stamen and pistil numbers. (**C**) Inferred optimal sex allocation strategies changed from predominantly male-biased toward highly female-biased for individuals with low and high resource status, respectively. Note that the slope of the trade-off lines is conceptual, because we do not know the linearity and the actual trade-off ratio relating female to male units of allocation.

## Discussion

### Estimation of male and female gain curves in a natural population of flowers

Our study provides a rare estimate of the shape of fitness gain curves for a natural plant population. It has hitherto been difficult to estimate the shape of fitness gain curves for several reasons, not least because most species do not display a sufficiently wide range of phenotypic variation in natural populations over which to estimate fitness. By removing some or all stamens of a sample of flowers of our study species *P. alpina*, we extended the already wide range of sex allocation represented in the population to include not only male and bisexual flowers, which occur naturally, but also fully female flowers. Because the range of sex allocation in the study population covered the full plane of allocations, we could estimate male and female gain curves relatively independently of one another^56^.

Our analyses indicate that the male fitness gain curve for *P. alpina* tended towards saturation in single-flowered individuals. Although there have been very few empirical estimates of the shape of the male gain curve for animal-pollinated plants^12,16^, those that do exist have also found evidence for saturating male gain curves^57–59^ (but see reference^60^). Saturating fitness gain curves are expected if pollen accumulation saturates on pollinators’ bodies^12^, or if pollen from a given individual is delivered to a small number of receptive stigmas, causing local mate competition^6,61,62^ or local sperm competition^15^. *Pulsatilla alpina* is pollinated almost exclusively by flies that cause substantial within-flower self-pollination^44^ and that disperse pollen among individuals over short distances (mean = 3.16 m)^63^, conditions that should give rise to local mate competition and thus to saturating male fitness gain curves.

In contrast with the tendency towards saturation for male gain curve, the female gain curve for single-flowered individuals of *P. alpina* was strongly accelerating, likely as a result of sexual interference between male and female allocation^64^. The female gain curve in plants is often suspected to be saturating rather than accelerating, because competition among progeny dispersed into a limited seed shadow (‘local resource competition’) compromises the seed producer’s inclusive fitness^12,65,66^. While we did not measure possible effects of local resource competition on the female gain curve, we believe this possibility is negligible in *P. alpina* because the single-seeded fruits (achenes) are furnished with parachute-like structures that aid their dispersal by wind from elongated floral stalks^67,68^. Accordingly, although pollen is dispersed over very short distances, there is almost no genetic structure in populations of *P. alpina*^63^, consistent with a scenario of well-dispersed seeds^69^. The accelerating fitness gain curve estimated for *P. alpina* flowers likely derives from the fact that flowers with more pistils and/or fewer stamens have a lower within-flower selfing rate as a result of an increased separation between pistils and stamens within flowers^44^ and thus a lower rate of seed discounting^54^. This explanation is confirmed by analysis that ignores the effects of inbreeding depression, which predicted a slightly (albeit not statistically significant) saturating female gain curve for flowers rather than the accelerating curve predicted by analysis that incorporates inbreeding depression into estimates of floral contributions to fitness.

### A landscape approach for interpreting complex sex allocation strategies

We found that individuals of *P. alpina* occupy a large part of the area on the male/female allocation plane, indicating that sex allocation is clearly not constrained to lie on a single straightforward trade-off diagonal assumed by sex-allocation theory (see Figure 1). Such a scenario is likely to be common in perennial plants and animals whose resource status varies with size or for other, more cryptic, reasons^15,70^. A corollary of our finding here is that the female component of reproductive success in *P. alpina* likely depends not only on female allocation, but also on male allocation in ways that go beyond linear sex-allocation trade-offs (and vice versa for male reproductive success). Our results indicate that the selfing rate and a flower’s contribution to female reproductive success was indeed a function not only of the number of pistils, but also of the number of stamens for a given pistil number^44^. Classic sex-allocation theory is unable to account for such patterns in a straightforward manner, though indirect effects of allocation to one sex on fitness through the other may be incorporated in a framework of sexual selection^71^. Indeed, although we found that the contribution to male reproductive success in *P. alpina* was independent of the number of pistils in single-flowered individuals, a previous study on the same species, but focusing only on bisexual flowers, found evidence for a cross-sex effect of this sort, consistent with the within-flower sexual conflict^45^.

To appreciate the value of mapping fitness components on a two-dimensional landscape of empirically measurable components of sex allocation when resource status obscures potentially underlying trade-offs, consider the diagonal lines traced on the fitness maps inferred for *P. alpina* in Figure 4A-C. Each of the lines represents a hypothetical linear sex-allocation trade-off, and a walk over the fitness landscape following these paths (Figure 5) illustrates how the shape of fitness gain curves may vary with plant resource status (i.e., small, medium and large). This depiction suggests that small individuals of *P. alpina* should allocate most of their reproductive resources to their male function, while larger individuals (with more resources) should allocate substantially to both male and female functions (Figure 5). With growth, individuals should thus shift from an all-male to a hermaphroditic allocation strategy, i.e., they should display a type of gender diphasy^21,36,72^, as indeed observed in wild populations of *P. alpina*^43^ and many other perennial plants^73–75^ and animals^76^.

The existence of two fitness peaks on the sex allocation landscape for *P. alpina* flowers (Figure 4) also points to andromonoecy as a successful strategy, with some flowers adopting a fully male and others a bisexual strategy. Again, this pattern corresponds qualitatively to the strategy adopted by *P. alpina* (Figure S3). By abandoning their female function and producing fully male flowers, individuals of *P. alpina* may capitalize on opportunities for mating in the early season when most of the other bisexual protogynous flowers are in their female stage^45^. It is plausible that individuals with substantial reproductive resources in general might set aside as much as they can to produce bisexual flowers and use whatever remains for one or more male flowers at the beginning of the season (note that all *P. alpina* floral buds are formed below ground at the end of the previous growing season^77^). In andromonoecious species of *Solanum*, for instance, there is a strong correlation between the size of the fruit and the fraction of male flowers, supporting the notion that male flowers in andromonoecious species in general may often serve to balance sex allocation in this way^78^.

Finally, the topography of the fitness landscape provides a clear indication of a saturating (rather than the otherwise accelerating) female fitness curve for flowers of *P. alpina* from which stamens had experimentally been removed, i.e., for a phenotype that does not occur in nature (because *P. alpina* flowers always bear stamens; Figures 4A and S4). *Pulsatilla alpina* stamens function not only to sire ovules but also to attract pollinators to flowers that do not produce nectar, so that the lower female reproductive success of flowers without stamens may have been a result of pollinator limitation – though this conjecture needs testing. Nonetheless, this possibility further exemplifies the utility of our approach in being able to reveal cross-sex effects when they exist.

The function of stamens as both the source of pollen for siring offspring as well as a reward for pollinators might further help to explain why the flowers of *P. alpina* always produce stamens and thus why the species is andromonoecious (with unisexual male and bisexual flowers) and not monoecious (with unisexual female and male flowers), likely resembling the cases of andromonoecy in *Solanum*^79–81^.

We have interpreted strategies at the floral level using a fitness landscape inferred from a population of mostly single-flowered individuals in a species that often produces multiple flowers. This needs to be borne in mind when interpreting a multi-flower strategy. First, inferences of positive marginal effects on fitness through the production of male flowers will need to be tempered if the flowers negatively affect the fitness contribution of other flowers with a female function, e.g., through increased geitonogamous selfing^82^ (but see^25^). This seems an unlikely scenario in *P. alpina*, because its male flowers tend to be open at the beginning of the flowering season and thus often function alone even for individuals that produce bisexual flowers^43^. Second, the inference of an accelerating fitness gain curve for the female function of individuals with only one flower would be misleading for individuals that disperse seeds from many flowers into a small seed shadow that then suffer from local resource competition^12,65,66^. As discussed above, we believe that local resource competition is unlikely to be important in *P. alpina*^67,68^. Finally, the fitness landscape we have inferred for single-flowered *P. alpina* individuals might seem to point to androdioecy (with distinct male and hermaphrodite individuals), as much as to andromonoecy. But multi-flowered males would produce their male flowers over the course of the reproductive season, yet the andromonoecious strategy of *P. alpina* involves producing one or a few male flowers only at the beginning of the season when the species’ strong protogyny ensures high mating opportunities^43,45^. It is well established that androdioecy is difficult to maintain when males must compete with hermaphrodites for siring opportunities, especially in partially selfing species^83,84^, as is the case of *P. alpina*^44^.

## Conclusion

Sex allocation theory has been successful in helping to explain variation in sex ratios in dioecious species, particularly animals, but its quantitative application to explain the complexity of sex-allocation strategies in hermaphrodites has had limited success. Our study has overcome key operational difficulties by successfully estimating both the male and female contributions to fitness by floral modules across their full potential range of allocations. It also showcases a simple but potentially useful approach for interpreting complex patterns of sex allocation in terms of the shape of a fitness landscape defined by independent measures of male and female allocation on orthogonal axes. This approach may provide an empirically accessible rescue-line for a body of theory that has been increasingly criticized for being too difficult to apply to the messy world of hermaphroditic reproduction^12,15,56,85,86^.

## Resource availability

### Lead contact

Further information and requests for resources should be directed to the lead contact, Kai-hsiu Chen.

### Materials availability

There are no new materials associated with this paper.

### Data and code availability

Data and codes used in this study can be accessed in Zenodo (DOI: 10.5281/zenodo.14289403).

## Acknowledgments

We thank Canton of Vaud, Commune of Bex, for access to field sites, N. Szijarto for help in field, D. Savova-Bianchi for help with data collection, and the University of Lausanne and the Swiss National Science Foundations (grant 310030_185196) for funding. We thank D. Charlesworth, E. Charnov, C. Mullon, and T. Lesaffre for valuable comments on a previous version of the manuscript.

## Statement of authorship

KHC and JRP designed the project and wrote the manuscript. KHC collected the data and led the data analysis, with input from JRP.

## Declaration of interests

The authors declare no competing interests.

## Materials and Methods

### Experimental model and study participant details

We studied a population of *Pulsatilla alpina* (L.) Delarbre (Ranunculaceae) at Solalex in the pre-Alps of Vaud canton, Switzerland (‘Population S1+’; latitude: 46°17′42″N, longitude: 7°09′09″E; elevation: 1758 a.s.l.) in the flowering season of 2022. The species grows in sub-alpine to alpine habitats in central Europe, with longevity likely exceeding 30 years^87^. Each spring, several vegetative and/or reproductive shoots emerge from a rhizome soon after the snowmelt, with single flowers on separate flowering shoots. Small plants often produce a single male or bisexual flower, while larger plants could produce up to about 20 usually bisexual flowers, i.e., the species displays gender diphasy as a result of size-dependent sex allocation^43^. Bisexual flowers have a similar number of stamens to that of male flowers and up to 400 single-ovulated pistils (Figure 1)^45^. Flowers are predominantly visited by flies^68^. Ripe fruits (technically achenes) with elongated pappus hairs are dispersed by wind in early autumn^67^. *Pulsatilla alpina* is self-compatible and has a selfing rate of about 0.47 (95% *CI*: 0.38 – 0.54; based on 1,000 bootstraps), with the rate of selfing being a positive function of within-flower male allocation^44^. Inbreeding depression was estimated to be 0.95 (0.92 – 0.96 under the 95% *CI* of mean selfing rate)^44^ based on comparisons of inbreeding coefficients (F_IS_) between adults and seed progeny in the study population^52^. As in most studies of inbreeding depression in plants, our estimate of inbreeding depression ignores the effect of inbreeding on seed development from the zygote and is thus somewhat conservative. The study population comprised a total of 104 individuals producing a single flower and 31 individuals producing more than one flower in 2022 (*N* = 135 flowering individuals, and a total of 175 flowers; Figure S1). The population was situated on an open slope of sub-alpine grassland that we fenced to exclude the most predominant herbivores, i.e., cattle and other browsers. We removed all floral buds from the few unsampled individuals outside the plot at the beginning of the flowering season to prevent them from siring progeny in the plot.

### Method details

#### Flowering phenology

We recorded the location and flowering state of all individuals in the population every three or four days from late May to late June, 2022 (in total twelve census times), noting the number and sexual phenotype of their flowers. For each flower and sampling date, we recorded each flower’s sexual stage in terms of seven and five ordinal categories for bisexual and male flowers, respectively (detailed description of the categories in Chen & Pannell^45^). These categories allowed us to manipulate flowers at the same developmental stage, i.e., before anther dehiscence.

#### Manipulation of floral sex allocation

Although flowers of *P. alpina* vary widely in their sex allocation, natural variation does not include flowers with few or no stamens. To estimate the potential fitness contributions of female or nearly fully female flowers, and thus to evaluate the full two-dimensional fitness landscape, we removed stamens of a sample of flowers at times throughout the flowering season, as has been done in other studies^32,56,88^. Prior to the stamen removal manipulations, 46 and 129 flowers were male and bisexual, respectively (Figure S1). At each observation time point, we randomly selected about a quarter of the bisexual flowers in their early female stage and used tweezers to remove 100% or 50% of their stamens. Similarly, we removed 50% of the stamens of about a quarter of male flowers in their early male stage^45^. Stamen removal did not alter the flowering duration of the flowers^45^, so its effect on components of fitness manifested only through changes in the number of stamens. Because stamen-removal occurred before anthers opened, there was no accidental intra-flower selfing.

#### Estimates of floral sex allocation

We quantified absolute floral sex allocation to the two sex functions as the number of stamens and pistils produced by each flower. For male allocation, we photographed all bisexual and male flowers at the late female stage and the mid_-_male stage (detailed description of the stages in Chen & Pannell^45^), respectively, and later counted the number of stamens based on the photographs calibrated against measures for a sample of 15 fresh flowers^43^. Three weeks after flowering ended, flowers with developing achenes were enclosed in individual paper bags to prevent seed dispersal, and seeds were later collected for counting. We quantified female allocation by counting the total number of achenes in each flower; each pistil, which contains a single ovule, develops into an achene and remains on the floral receptacle until the end of the season, whether fertilized or not^68^.

#### Paternity assignment and estimates of floral selfing rates

We first counted the number of mature seeds from a total of 22,612 achenes belonging to 104 seed families (19 and 6 seed families from 129 bisexual flowers were aborted or missing, respectively). Each seed family produced 90.5 ± 55.4 mature seeds (*N* = 104; mean ± SD). The seeds were well mixed and randomly selected from each seed family for genotyping. We extracted DNA (BioSprint 96 DNA Plant Kit), conducted PCR (HotStarTaq Master Mix Kit), and aimed to assign paternity to each of ten seeds sampled randomly from each flower in our sample, using variation at ten microsatellite loci (manually checked in GeneMapper v 6. 0) and the software Cervus v 3.0.7 set with a confidence level of 80% and an error rate of 0.018 (see details of the genetic markers in Chen & Pannell^45^). The ten genetic markers were highly polymorphic in the study population, with a mean of 8.7 alleles per locus and a mean polymorphic information content (PIC) of 0.65. We included all the flowering individuals (i.e., all single- and multiple-flowered individuals) as potential sires in the paternity analysis, none of which had an identical genotype. Furthermore, the average kinship coefficient F_r_ was -0.0001 over all the flowering individuals^63^, reassuring the accuracy of our paternity analysis^89^. In total, 892 of 1054 sampled seeds could be genotyped for at least five loci and used for paternity analysis (PCR was unsuccessful for the remaining seeds and were not included). The selfing rate was calculated by dividing the number of selfed seeds by the total number of paternity-assigned seeds for each flower.

#### Estimates of female reproductive success

We calculated components of reproductive success assuming both the estimated value of inbreeding depression (δ = 0.95), as well as assuming no inbreeding depression (δ = 0), with female reproductive success (RS_F_) computed as the inferred number of mature outcrossed seeds plus (1 – δ) times the inferred number of mature seeds produced by selfing, based on the number of mature seeds (N_T_) and the selfing rate (*S*) estimated for each bisexual flower with the following formula: RS_F_ = N_T_ × (1 - S) + N_T_ × S (1-*δ*).

#### Estimates of male reproductive success

We calculated male reproductive success for each flower as the number of outcrossed seeds sired on other individuals in the population (seed families from all the flowering individuals; *N* = 104) plus the number of seeds sired by selfing multiplied by (1 – δ), again for both δ = 0.95 and 0.0, as described before. Because we successfully assigned paternity for about eight seeds for all flowers (total *N* = 854), irrespective of the total number of seeds produced by the flower, we estimated the male reproductive success of a given potential sire by multiplying the fraction of the seeds in the flower it sired by the total number of seeds in that flower (mean correlated paternity^90^ was 0.19 ± 0.27 based on 88 seed families that had at least two outcrossed seeds). It is worth noting that although it would always be useful to increase the sample size of seeds from one seed family to increase the precision of the estimates of the selfing rate and paternity share, the power of our analysis lies in the replication of phenotypes varying in their sex allocation in the population (*N* = 87 individuals in total as the experimental units in the following analyses). A supplementary analysis using five seeds subsampled per family (total *N* = 497 seeds; all seeds were used if the family produced less than five seeds) yielded an almost identical fitness landscape, indicating that our fitness inferences based on estimated selfing rates and paternity shares are robust (Figure S5).

#### Quantification and statistical analysis

All the statistical analyses were conducted in R studio (v2024.12.1+563) with R (4.3.1)^91^. To investigate the relationship between components of fitness to floral sex allocation, we focused on female, male, and total reproductive success for all individuals that had only one flower (*N* = 87; 17 individuals with its flower aborted or missing were excluded). Because we wished to measure the total strength of selection (direct and indirect effects via correlation with other traits) on female and male sex allocation^41^, we excluded floral traits from our model, e.g., stalk height, petal length, and flowering date. Nonetheless, including the floral traits as covariates for estimating the strength of direct selection on sex allocation yielded qualitatively highly consistent results (see Figure S6), presumably because reproductive success is primarily determined by sex allocation in *P. alpina*. Moreover, a previous study^45^ found that stamen removal did not alter the flowering duration of *P. alpina* flowers, with the treatment effect on components of fitness manifested only through changes in stamen number.

We related prospective reproductive success to sex allocation using nonlinear univariate least square models with a Gaussian error distribution (*nls* function in R *stats*^91^) to evaluate the shape of fitness gain curves for female and male functions at the flower level, assuming both δ = 0 or δ = 0.95. The gain curve *y* was modelled as *y* = *ax^b^*, where *y* is the female or male reproductive success relative to the mean of that sex function for single-flowered individuals, *x* is the number of pistils or stamens for the female and male functions, respectively, *a* is a constant, and *b* < 1 and *b* > 1 correspond to a saturating or accelerating dependence of fitness on sex allocation^32,92^. *P-*values were calculated using a one-sample *t*-test.

We also used conventional selection gradient analysis with multivariate regression to evaluate the dependency of reproductive success on both the female and male components of allocation (i.e., as two phenotypic traits^93^), including second-order polynomial and interaction terms under the two scenarios of inbreeding depression at the flower level^53^. We fitted relative female, male, and total reproductive success (for both scenarios of inbreeding depression) as a response variable in six separate models, each with a Gaussian error distribution (*lm* function in R *stats*^91^). We standardized pistil number and stamen number to a mean of zero and a standard deviation of one, setting linear, quadratic, and interaction terms for the two traits in each model to evaluate linear and non-linear (i.e., quadratic and correlational) selection gradients on female and male allocation^53,94^. For all quadratic gradients, we multiplied the regression coefficients by two to obtain the correct estimate of stabilizing or disruptive selection^95^. *P*-values of the estimates were calculated using *t*-tests in six multivariate regression models.

Finally, we used nonparametric regression with smoothing functions to characterize the fitness landscapes for reproductive success in terms of female and male allocations under the two inbreeding depression scenarios^55,96^. Here, we used generalized additive models (*gam* function in R package *mgcv*^97^) for the dependency of components of reproductive success (as the numbers of seeds produced and/or sired) on female and male allocations on pistil and stamen number, assuming a Poisson error distribution; assuming negative-binomial or zero-inflated Poisson distributions yielded qualitatively identical results but explained less variance (Figure S6). We applied the *gam.gradients* function in the *R* package *GSG* to extract standardized linear, quadratic, and correlational selection gradients from the fitted models and calculated the standard errors and *P* values based on 1,000 bootstraps^55^. To plot the fitness surface, we used a smoothing term with thin plate splines^97^.

## Notes

### Competing Interest Statement

The authors have declared no competing interest.

### Summary of Updates

This version of the manuscript has been revised to update the presentation of the results.

## References

1. Charnov, E.L. (1982). The Theory of Sex Allocation 1st ed. (Princeton University Press).

2. Charlesworth, D., and Charlesworth, B. (1981). Allocation of resources to male and female functions in hermaphrodites. Biol. J. Linn. Soc. 15, 57–74. 10.1111/j.1095-8312.1981.tb00748.x.

3. West, S. (2009). Sex Allocation (Princeton University Press) 10.1515/9781400832019.

4. Hardy, I.C.W. (2002). Sex Ratios: Concepts and Research Methods (Cambridge University Press) 10.1017/CBO9780511542053.

5. Charnov, E.L., and Bull, J. (1977). When is sex environmentally determined? Nature 266, 828–830. 10.1038/266828a0.

6. Lloyd, D.G. (1984). Gender allocation in outcrossing cosexual plants. In Perspectives on Plant Population Ecology, R. Dirzo and J. Sarukhán, eds. (Sinauer Associates), pp. 277– 303.

7. Warner, R.R. (1988). Sex change and the size-advantage model. Trends Ecol. Evol. 3, 133–136. 10.1016/0169-5347(88)90176-0.

8. Klinkhamer, P., de Jong, T., and Metz, H. (1997). Sex and size in cosexual plants. Trends Ecol. Evol. 12, 260–265. 10.1016/S0169-5347(97)01078-1.

9. Ghiselin, M.T. (1969). The evolution of hermaphroditism among animals. Q. Rev. Biol. 44, 189–208. 10.1086/406066.

10. Orzack, S.H. (2009). Using sex ratios: the past and the future. In Sex Ratios, I. C. W. Hardy, ed. (Cambridge University Press), pp. 383–398. 10.1017/cbo9780511542053.020.

11. Brunet, J. (1992). Sex allocation in hermaphroditic plants. Trends Ecol. Evol. 7, 79–84. 10.1016/0169-5347(92)90245-7.

12. Campbell, D.R. (2000). Experimental tests of sex-allocation theory in plants. Trends Ecol. Evol. 15, 227–232. 10.1016/S0169-5347(00)01872-3.

13. Charlesworth, D., and Morgan, M.T. (1991). Allocation of resources to sex functions in flowering plants. Philos. Trans. R. Soc. London. Ser. B Biol. Sci. 332, 91–102. 10.1098/rstb.1991.0037.

14. Charnov, E.L., Smith, J.M., and Bull, J. (1976). Why be an hermaphrodite? Nature 263, 125–126. 10.1038/263125a0.

15. Schärer, L. (2009). Tests of sex allocation theory in simultaneously hermaphroditic animals. Evolution. 63, 1377–1405. 10.1111/J.1558-5646.2009.00669.X.

16. de Jong, T.J., and Klinkhamer, P.G.L. (2005). Evolutionary Ecology of Plant Reproductive Strategies (Cambridge University Press).

17. Munday, P.L., Buston, P.M., and Warner, R.R. (2006). Diversity and flexibility of sex-change strategies in animals. Trends Ecol. Evol. 21, 89–95. 10.1016/j.tree.2005.10.020.

18. Sinclair, J.P., Emlen, J., and Freeman, D.C. (2012). Biased sex ratios in plants: Theory and trends. Bot. Rev. 78, 63–86. 10.1007/S12229-011-9065-0/FIGURES/2.

19. Ashman, T.L. (2003). Constraints on the evolution of males and sexual dimorphism: field estimates of genetic architecture of reproductive traits in three populations of gynodioecious *Fragaria virginiana*. Evolution. 57, 2012–2025. 10.1111/J.0014-3820.2003.TB00381.X.

20. Mazer, S.J., Delesalle, V.A., and Paz, H. (2007). Evolution of mating system and the genetic covariance between male and female investments in *Clarkia* (Onagraceae): selfing opposes the evolution of trade-offs. Evolution. 61, 83–98. 10.1111/J.1558-5646.2007.00007.X.

21. Zhang, D.Y., and Jiang, X.H. (2002). Size-dependent resource allocation and sex allocation in herbaceous perennial plants. J. Evol. Biol. 15, 74–83. 10.1046/j.1420-9101.2002.00369.x.

22. Sakai, S. (2000). Biased sex allocation in hermaphroditic plants. J. Plant Res. 113, 335–342. 10.1007/PL00013937.

23. Dorken, M.E., and Van Drunen, W.E. (2018). Life-history trade-offs promote the evolution of dioecy. J. Evol. Biol. 31, 1405–1412. 10.1111/jeb.13335.

24. Seger, J., and Eckhart, V.M. (1996). Evolution of sexual systems and sex allocation in plants when growth and reproduction overlap. Proc. R. Soc. B Biol. Sci. 263, 833–841. 10.1098/rspb.1996.0123.

25. Tomaszewski, C.E., Kulbaba, M.W., and Harder, L.D. (2018). Mating consequences of contrasting hermaphroditic plant sexual systems. Evolution. 72, 2114–2128. 10.1111/evo.13572.

26. Oddou-Muratorio, S., Bontemps, A., Gauzere, J., and Klein, E.K. (2024). Interplay between fecundity, sexual and growth selection on the spring phenology of European beech (*Fagus sylvatica* L.). Peer Community J. 4, e27. 10.24072/pcjournal.396.

27. Kwok, A., and Dorken, M.E. (2022). Sexual selection on male but not female function in monoecious and dioecious populations of broadleaf arrowhead (*Sagittaria latifolia*). Proc. R. Soc. B Biol. Sci. 289, 20220919. 10.1098/rspb.2022.0919.

28. Angeloni, F., Ouborg, N.J., and Leimu, R. (2011). Meta-analysis on the association of population size and life history with inbreeding depression in plants. Biol. Conserv. 144, 35–43. 10.1016/j.biocon.2010.08.016.

29. de Jong, T.J., Klinkhamer, P.G.L., and Rademaker, M.C.J. (1999). How geitonogamous selfing affects sex allocation in hermaphrodite plants. J. Evol. Biol. 12, 166–176. 10.1046/J.1420-9101.1999.00001.X.

30. Nakahara, T., Fukano, Y., Hirota, S.K., and Yahara, T. (2018). Size advantage for male function and size-dependent sex allocation in *Ambrosia artemisiifolia*, a wind-pollinated plant. Ecol. Evol. 8, 1159–1170. 10.1002/ece3.3722.

31. Tonnabel, J., David, P., Klein, E.K., and Pannell, J.R. (2019). Sex-specific selection on plant architecture through “budget” and “direct” effects in experimental populations of the wind-pollinated herb, *Mercurialis annua*. Evolution. 73, 897–912. 10.1111/EVO.13714.

32. Aljiboury, A.A., and Friedman, J. (2022). Mating and fitness consequences of variation in male allocation in a wind-pollinated plant. Evolution. 76, 1762–1775. 10.1111/evo.14544.

33. Diggle, P.K. (2003). Architectural effects on floral form and function: a review. In Deep Morphology: Toward a Renaissance of Morphology in Plant Systematics, T. F. Stuessy, V. Mayer, and E. Horandl, eds. (Koeltz), pp. 63–80.

34. Harder, L.D., and Prusinkiewicz, P. (2013). The interplay between inflorescence development and function as the crucible of architectural diversity. Ann. Bot. 112, 1477– 1493. 10.1093/aob/mcs252.

35. Cardoso, J.C.F., Viana, M.L., Matias, R., Furtado, M.T., Caetano, A.P. de S., Consolaro, H., and Brito, V.L.G. de (2018). Towards a unified terminology for angiosperm reproductive systems. Acta Bot. Brasilica 32, 329–348. 10.1590/0102-33062018ABB0124.

36. Schlessman, M.A. (1988). Gender diphasy (“sex choice”). In Plant Reproductive Ecology: Patterns and Strategies, J. Lovett-Doust and L. Lovett-Doust, eds. (Oxford University Press), pp. 139–153.

37. Lloyd, D.G., and Webb, C.J. (1986). The avoidance of interference between the presentation of pollen and stigmas in angiosperms I. Dichogamy. New Zeal. J. Bot. 24, 135–162. 10.1080/0028825X.1986.10409725.

38. Brunet, J., and Charlesworth, D. (1995). Floral sex allocation in sequentially blooming plants. Evolution. 49, 70–79. 10.2307/2410293.

39. de Jong, T.J., Shmida, A., and Thuijsman, F. (2008). Sex allocation in plants and the evolution of monoecy. Evol. Ecol. Res. 10, 1087–1109.

40. Bataillon, T., Ezard, T.H.G., Kopp, M., and Masel, J. (2022). Genetics of adaptation and fitness landscapes: from toy models to testable quantitative predictions. Evolution. 76, 1104–1107. 10.1111/EVO.14477.

41. Brodie, E.D., Moore, A.J., and Janzen, F.J. (1995). Visualizing and quantifying natural selection. Trends Ecol. Evol. 10, 313–318. 10.1016/S0169-5347(00)89117-X.

42. Saeki, Y., Tuda, M., and Crowley, P.H. (2014). Allocation tradeoffs and life histories: a conceptual and graphical framework. Oikos 123, 786–793. 10.1111/OIK.00956.

43. Chen, K.-H., and Pannell, J.R. (2023). Size-dependent sex allocation and the expression of andromonoecy in a protogynous perennial herb: both size and timing matter. bioRxiv, 2023.03.10.532080. 10.1101/2023.03.10.532080.

44. Chen, K.-H., and Pannell, J.R. (2024). Ignoring withinDflower selfDfertilization and inbreeding depression biases estimates of selection on floral traits in a perennial alpine herb. J. Ecol. 112, 2540–2551. 10.1111/1365-2745.14378.

45. Chen, K.-H., and Pannell, J.R. (2023). Unisexual flowers as a resolution to intralocus sexual conflict in hermaphrodites. Proc. R. Soc. B Biol. Sci. 290, 20232137. 10.1098/rspb.2023.2137.

46. Burd, and Callahan (2000). What does the male function hypothesis claim? J. Evol. Biol. 13, 735–742. 10.1046/j.1420-9101.2000.00220.x.

47. Whitehead, M.R., Lanfear, R., Mitchell, R.J., and Karron, J.D. (2018). Plant mating systems often vary widely among populations. Front. Ecol. Evol. 6, 1–9. 10.3389/fevo.2018.00038.

48. Hou, M., Opedal, Ø.H., and Zhao, Z.G. (2024). Sexually concordant selection on floral traits despite greater opportunity for selection through male fitness. New Phytol. 241, 926– 936. 10.1111/nph.19370.

49. Briscoe Runquist, R.D., Geber, M.A., Pickett-Leonard, M., and Moeller, D.A. (2017). Mating system evolution under strong pollen limitation: evidence of disruptive selection through male and female fitness in *Clarkia xantiana*. Am. Nat. 189, 549–563. 10.1086/691192.

50. Austen, E.J., and Weis, A.E. (2016). Estimating selection through male fitness: three complementary methods illuminate the nature and causes of selection on flowering time. Proc. R. Soc. B Biol. Sci. 283, 20152635. 10.1098/rspb.2015.2635.

51. Kulbaba, M.W., and Shaw, R.G. (2021). Lifetime fitness through female and male function: Influences of genetically effective population size and density. Am. Nat. 197, 434–447. 10.1086/713067.

52. Ritland, K. (1990). Inferences about inbreeding depression based on changes of the inbreeding coefficient. Evolution. 44, 1230–1241. 10.1111/J.1558-5646.1990.TB05227.X.

53. Lande, R., and Arnold, S.J. (1983). The measurement of selection on correlated characters. Evolution. 37, 1210–1226. 10.2307/2408842.

54. Lloyd, D.G. (1992). Self- and cross-fertilization in plants. II. The selection of self-fertilization. Int. J. Plant Sci. 153, 370–380. 10.1086/297041.

55. Morrissey, M.B., and Sakrejda, K. (2013). Unification of regression-based methods for the analysis of natural selection. Evolution. 67, 2094–2100. 10.1111/EVO.12077.

56. Emms, S.K. (1993). On measuring fitness gain curves in plants. Ecology 74, 1750–1756. 10.2307/1939933.

57. Campbell, D.R. (1998). Variation in lifetime male fitness in Ipomopsis aggregata: tests of sex allocation theory. Am. Nat. 152, 338–353. 10.1086/286173.

58. Rademaker, M.C.J., and De Jong, T.J. (1998). Effects of flower number on estimated pollen transfer in natural populations of three hermaphroditic species: an experiment with fluorescent dye. J. Evol. Biol. 11, 623–641. 10.1046/j.1420-9101.1998.11050623.x.

59. Rosas, F., and Domínguez, C.A. (2009). Male sterility, fitness gain curves and the evolution of gender specialization from distyly in *Erythroxylum havanense*. J. Evol. Biol. 22, 50–59. 10.1111/j.1420-9101.2008.01618.x.

60. Perry, L.E., and Dorken, M.E. (2011). The evolution of males: support for predictions from sex allocation theory using mating arrays of *Sagittaria latifolia* (Alimataceae). Evolution. 65, 2782–2791. 10.1111/J.1558-5646.2011.01344.X.

61. Hamilton, W.D. (1967). Extraordinary sex ratios. Science. 156, 477–488. 10.1126/science.156.3774.477.

62. Harder, L.D., and Johnson, S.D. (2025). Pollination efficiency and the evolution of sex allocation – diminishing returns matter. New Phytol. 246, 39–46. 10.1111/NPH.20389.

63. Chen, K.-H., and Pannell, J.R. (2024). Pollen dispersal distance is determined by phenology and ancillary traits but not floral gender in an andromonoecious, fly-pollinated alpine herb. Alp. Bot. 134, 69–79. 10.1007/s00035-024-00313-z.

64. Barrett, S.C.H. (2002). Sexual interference of the floral kind. Heredity. 88, 154–159. 10.1038/sj.hdy.6800020.

65. Takahashi, K., Makino, T.T., and Sakai, S. (2005). Effects of sib-competition on female reproductive success in *Salvia lutescens* Koidz. var. crenata. Evol. Ecol. Res. 7, 1201– 1212.

66. Rademaker, M.C.J., and De Jong, T.J. (1999). The shape of the female fitness curve for *Cynoglossum officinale*: quantifying seed dispersal and seedling survival in the field. Plant Biol. 1, 351–356. 10.1111/j.1438-8677.1999.tb00263.x.

67. Vittoz, P., and Engler, R. (2007). Seed dispersal distances: a typology based on dispersal modes and plant traits. Bot. Helv. 117, 109–124. 10.1007/s00035-007-0797-8.

68. Chen, K.-H., and Pannell, J.R. (2022). Disruptive selection via pollinators and seed predators on the height of flowers on a wind-dispersed alpine herb. Am. J. Bot. 109, 1717–1729. 10.1002/AJB2.16073.

69. Vekemans, X., and Hardy, O.J. (2004). New insights from fine-scale spatial genetic structure analyses in plant populations. Mol. Ecol. 13, 921–935. 10.1046/J.1365-294X.2004.02076.X.

70. Sarkissian, T.S., Barrett, S.C.H., and Harder, L.D. (2001). Gender variation in *Sagittaria latifolia* (Alismataceae): is size all that matters? Ecology 82, 360–373. 10.1890/0012-9658(2001)082[0360:gvisla]2.0.co;2.

71. Anthes, N., David, P., Auld, J.R., Hoffer, J.N.A., Jarne, P., Koene, J.M., Kokko, H., Lorenzi, M.C., Pélissié, B., Sprenger, D., et al. (2010). Synthesis Bateman gradients in hermaphrodites: an extended approach to quantify sexual selection. Am. Nat. 176, 249–263. 10.1086/655218/SUPPL_FILE/51575APB.PDF.

72. Freeman, D.C., Harper, K.T., and Charnov, E.L. (1980). Sex change in plants: old and new observations and new hypotheses. Oecologia 47, 222–232. 10.1007/BF00346825.

73. Niu, Y., Gong, Q., Peng, D., Sun, H., and Li, Z. (2017). Function of male and hermaphroditic flowers and size-dependent gender diphasy of *Lloydia oxycarpa* (Liliaceae) from Hengduan Mountains. Plant Divers. 39, 187–193. 10.1016/j.pld.2017.06.001.

74. Astuti, G., Pratesi, S., Carta, A., and Peruzzi, L. (2020). Male flowers in *Tulipa pumila* Moench (Liliaceae) potentially originate from gender diphasy. Plant Species Biol. 35, 130–137. 10.1111/1442-1984.12267.

75. Schlessman, M.A. (1991). Size, gender, and sex change in dwarf ginseng, *Panax trifolium* (Araliaceae). Oecologia 87, 588–595. 10.1007/BF00320425.

76. Baeza, J.A. (2007). Sex allocation in a simultaneously hermaphroditic marine shrimp. Evolution. 61, 2360–2373. 10.1111/J.1558-5646.2007.00199.X.

77. Jutila, H., Parisy, B., and Loehr, J. (2024). Influence of environmental and intrinsic factors on the flowering success and petal morphology of *Pulsatilla patens* and the hybrid *Pulsatilla patens* × *vernalis* in Finland. Plant Ecol. 225, 425–440. 10.1007/s11258-024-01400-1.

78. Miller, J.S., and Diggle, P.K. (2007). Correlated evolution of fruit size and sexual expression in andromonoecious *Solanum* sections *Acanthophora* and *Lasiocarpa* (Solanaceae). Am. J. Bot. 94, 1706–1715. 10.3732/AJB.94.10.1706.

79. Anderson, G.J., and Symon, D.E. (1989). Functional dioecy and andromonoecy in *Solanum*. Evolution. 43, 204–219. 10.1111/j.1558-5646.1989.tb04218.x.

80. Connolly, B.A., and Anderson, G.J. (2003). Functional significance of the androecium in staminate and hermaphroditic flowers of *Solanum carolinense* (Solanaceae). Plant Syst. Evol. 240, 235–243. 10.1007/s00606-003-0029-7.

81. Russell, A.L., Golden, R.E., Leonard, A.S., and Papaj, D.R. (2016). Bees learn preferences for plant species that offer only pollen as a reward. Behav. Ecol. 27, 731–740. 10.1093/BEHECO/ARV213.

82. Spalik, K. (1991). On evolution of andromonoecy and ‘overproduction’ of flowers: a resource allocation model. Biol. J. Linn. Soc. 42, 325–336. 10.1111/j.1095-8312.1991.tb00566.x.

83. Charlesworth, D. (1984). Androdioecy and the evolution of dioecy. Biol. J. Linn. Soc. 22, 333–348. 10.1111/J.1095-8312.1984.TB01683.X.

84. Pannell, J.R. (2002). The evolution and maintenance of androdioecy. Annu. Rev. Ecol. Syst. 33, 397–425. 10.1146/ANNUREV.ECOLSYS.33.010802.150419.

85. Thomson, J.D. (2006). Tactics for male reproductive success in plants: contrasting insights of sex allocation theory and pollen presentation theory. Integr. Comp. Biol. 46, 390–397. 10.1093/ICB/ICJ046.

86. Burd, M. (2024). Why the Shaw–Mohler equation works and when it doesn’t. Biol. Lett. 20, 20230499. 10.1098/rsbl.2023.0499.

87. Edelfeldt, S., Lindell, T., and Dahlgren, J.P. (2019). Age-independent adult mortality in a long-lived herb. Diversity 11, 187. 10.3390/D11100187.

88. Johnson, S.L., and Yund, P.O. (2009). Effects of fertilization distance on male gain curves in a free-spawning marine invertebrate: a combined empirical and theoretical approach. Evolution. 63, 3114–3123. 10.1111/J.1558-5646.2009.00784.X.

89. Marshall, T.C., Slate, J., Kruuk, L.E.B., and Pemberton, J.M. (1998). Statistical confidence for likelihood-based paternity inference in natural populations. Mol. Ecol. 7, 639–655. 10.1046/j.1365-294x.1998.00374.x.

90. Ritland, K. (1989). Correlated matings in the partial selfer *Mimulus guttatus*. Evolution. 43, 848–859. 10.1111/j.1558-5646.1989.tb05182.x.

91. R Core Team (2021). R: a language and environment for statistical computing.

92. Charnov, E.L. (1979). Simultaneous hermaphroditism and sexual selection. Proc. Natl. Acad. Sci. U. S. A. 76, 2480–2484. 10.1073/pnas.76.5.248.

93. Vallejo-Marín, M., and Rausher, M.D. (2007). Selection through female fitness helps to explain the maintenance of male flowers. Am. Nat. 169, 563–568. 10.1086/513112.

94. Matsumura, S., Arlinghaus, R., and Dieckmann, U. (2012). Standardizing selection strengths to study selection in the wild: a critical comparison and suggestions for the future. Bioscience 62, 1039–1054. 10.1525/BIO.2012.62.12.6.

95. Stinchcombe, J.R., Agrawal, A.F., Hohenlohe, P.A., Arnold, S.J., and Blows, M.W. (2008). Estimating nonlinear selection gradients using quadratic regression coefficients: double or nothing? Evolution. 62, 2435–2440. 10.1111/J.1558-5646.2008.00449.X.

96. Schluter, D., and Nychka, D. (1994). Exploring fitness surfaces. Am. Nat. 143, 597–616. 10.1086/285622.

97. Wood, S.N. (2003). Thin plate regression splines. J. R. Stat. Soc. Ser. B Stat. Methodol. 65, 95–114. 10.1111/1467-9868.00374.

