## Supplementary Information for "Mapping fitness landscapes to interpret sex allocation in hermaphrodites"

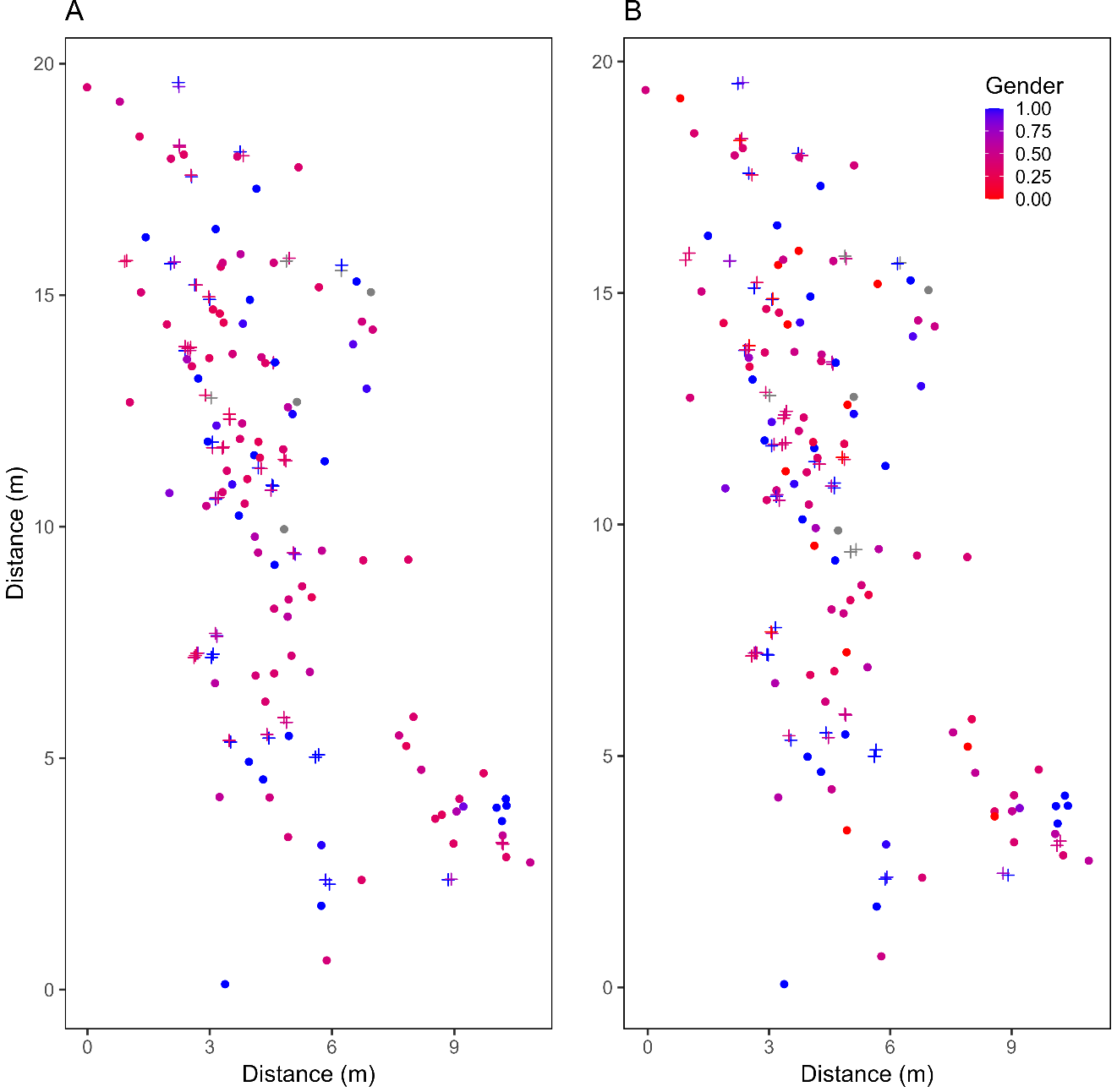

**Figure S1.** Position of the flowers in Population S1+ of *Pulsatilla alpina* and their relative male allocation in terms of gender (maleness) based on stamen and pistil numbers (**A**) before and (**B**) after stamen removal manipulation. Each circle and cross represents a flower from single-flowered (*N* = 104) and multiple-flowered individuals (*N* = 71 flowers from 31 individuals), respectively. Points are jittered to avoid overlapping. The grey points represent missing data (*N* = 8). Individuals were distributed randomly over space in terms of their functional gender both before (Moran index = -0.01, *P* = 0.62) and after (Moran index = -0.002, *P* = 0.75) manipulation; the spatial pattern of sex allocation was thus not further in analyses.

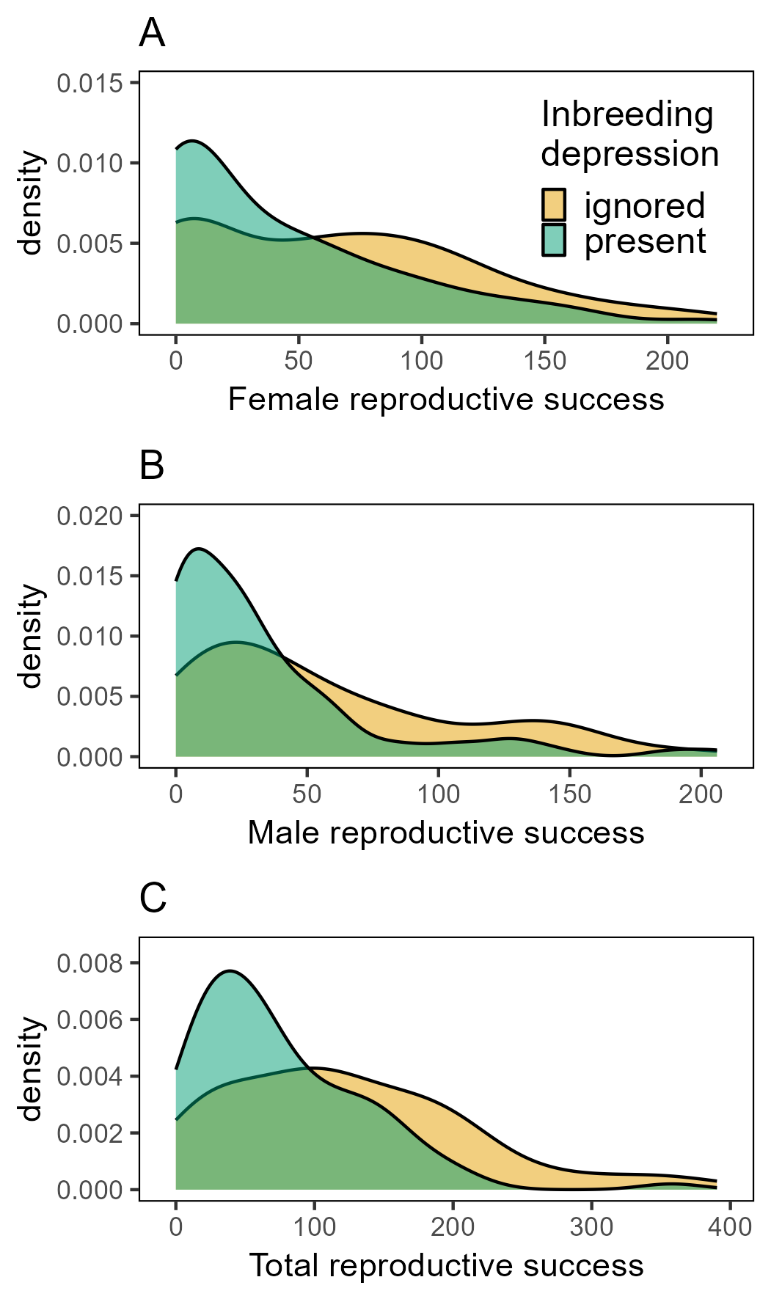

**Figure S2.** Density plots of inferred female (**A**), male (**B**), and total (**C**) reproductive success of single-flowered individuals of *Pulsatilla alpina* under two scenarios of inbreeding depression (*N* = 87). Reproductive success represents the actual number of seeds sired and/or produced. Scenarios of an inbreeding depression of zero and 0.95 are shown in yellow and green, respectively. Mean reproductive success of the single-flowered individuals did not differ between the two sexual functions for either scenario of inbreeding depression (*t*-test; when *δ* = 0, *t* = -0.94, *P* > 0.05; when *δ* = 0.95, *t* = -1.13, *P* > 0.05).

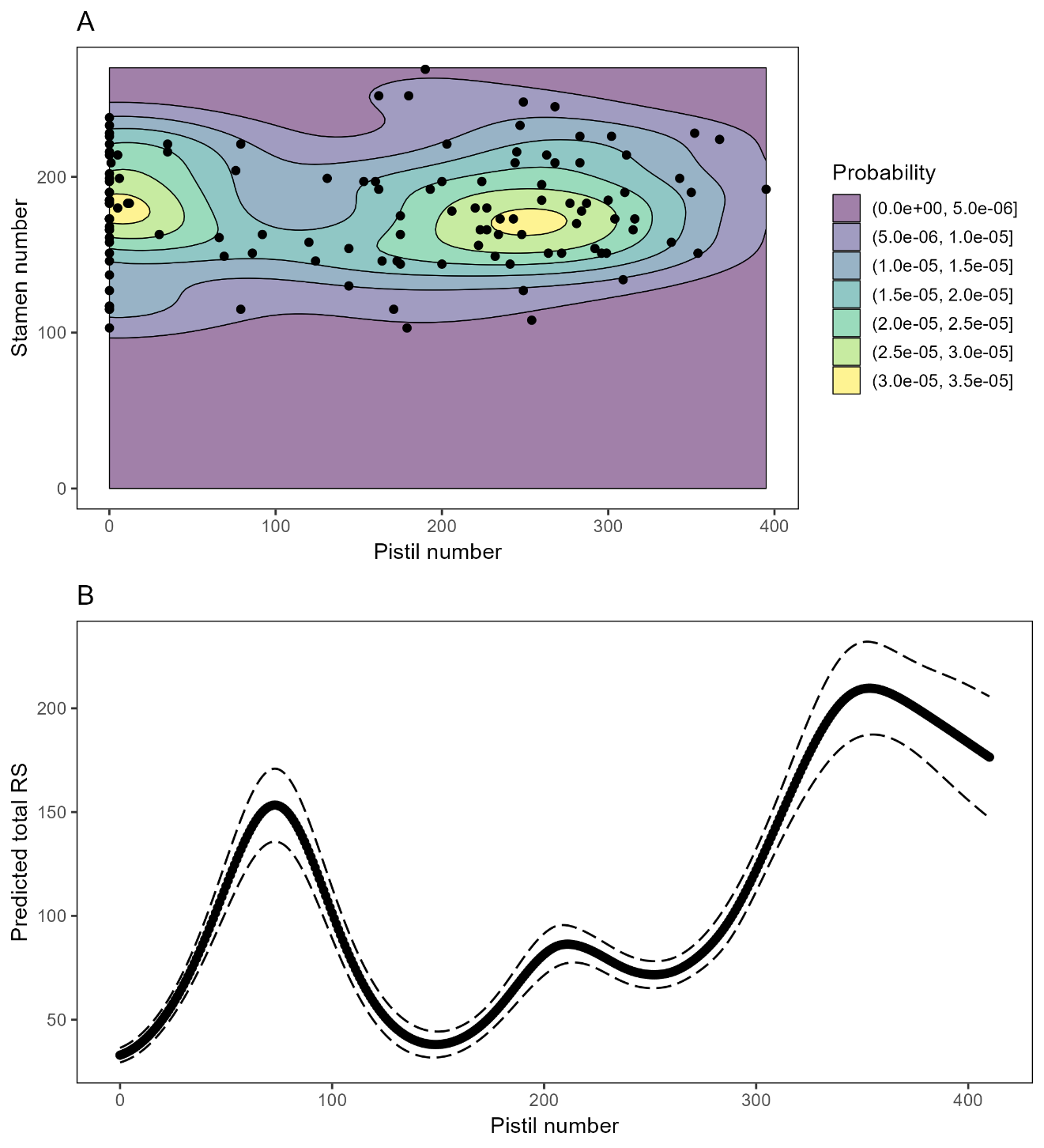

**Figure S3.** (**A**) Probability distribution of adopted sex allocation strategies by intact flowers of *Pulsatilla alpina* (*N* = 122; including intact flowers from both single and multiple-flowered individuals). Colour gradient from purple to yellow indicates low to high probability, respectively. Black points indicate the flowers. A supplementary analysis confirmed a positive albeit weak correlation (*r* = 0.11; Pearson correlation test; *t* = 3.43, *df* = 998, *P* = 0.0006) between the probability of a floral phenotype adopted by intact flowers and the projected total fitness of that phenotype (*δ* = 0.95) by randomly drawing 1000 phenotypes from the morphological space. This analysis indicates that the floral sex allocation strategy adopted by natural individuals qualitatively follows the topology of our fitness projection. (**B**) Projected total fitness (bold line) and its 95% confidence interval (dashed lines) along a linear path through the landscape for a scenario of 200 stamens and passing through the two fitness peaks in Figure **4E** in the main text. This path illustrates the significance of the two fitness peaks from the valleys.

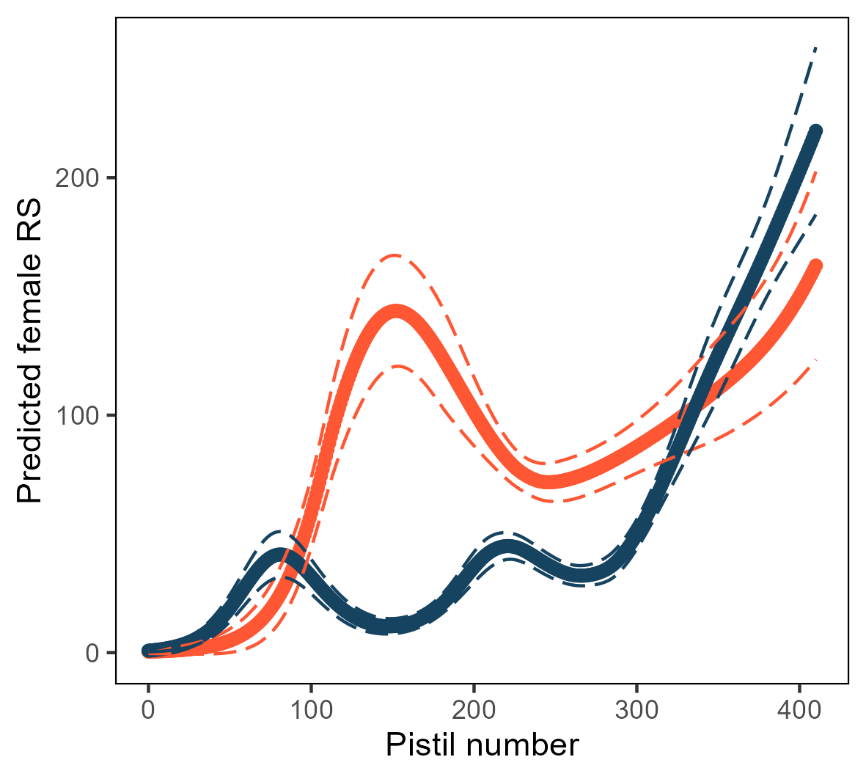

**Figure S4.** Contrasting female fitness gain curves for flowers with a fixed number of zero (red line) or 180 (blue line) stamens projected from the female fitness landscape presented in Figure **4A** when inbreeding depression was taken into account. Dashed lines indicate the 95% confidence intervals for the two scenarios. The unisexual female flowers created by experimental stamen removal likely had a saturating fitness return for their female function (red line). In contrast, the female fain curve was accelerating for flowers with a mean number of stamens (mean over all the intact flowers), resembling the patterns presented in Figures **2A** and **3A**.

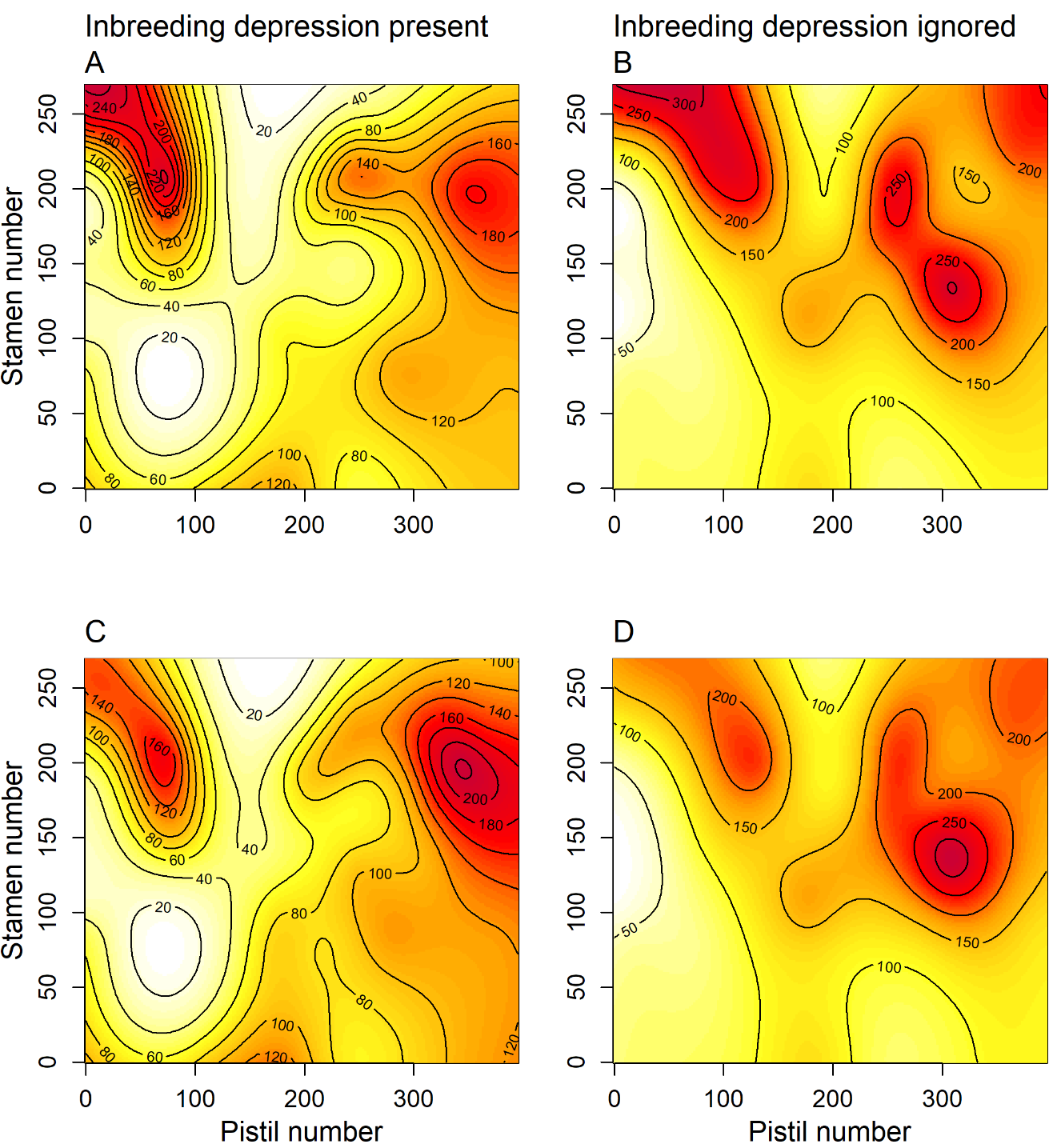

**Figure S5.** Plot comparing the fitness landscapes of total reproductive success for *Pulsatilla alpina* estimated by around half (58%) the sampling effort (**A** and **B**; subsampling 5 seeds genotyped per family; *N* = 497 seeds) and current sampling effort (**C** and **D**; *N* = 854 seeds) under the two inbreeding depression scenarios (related to STAR Methods). Note that the fitness landscapes remain largely the same, no matter which sampling effort is adopted. The predicted fitness was shown as the number of seeds.

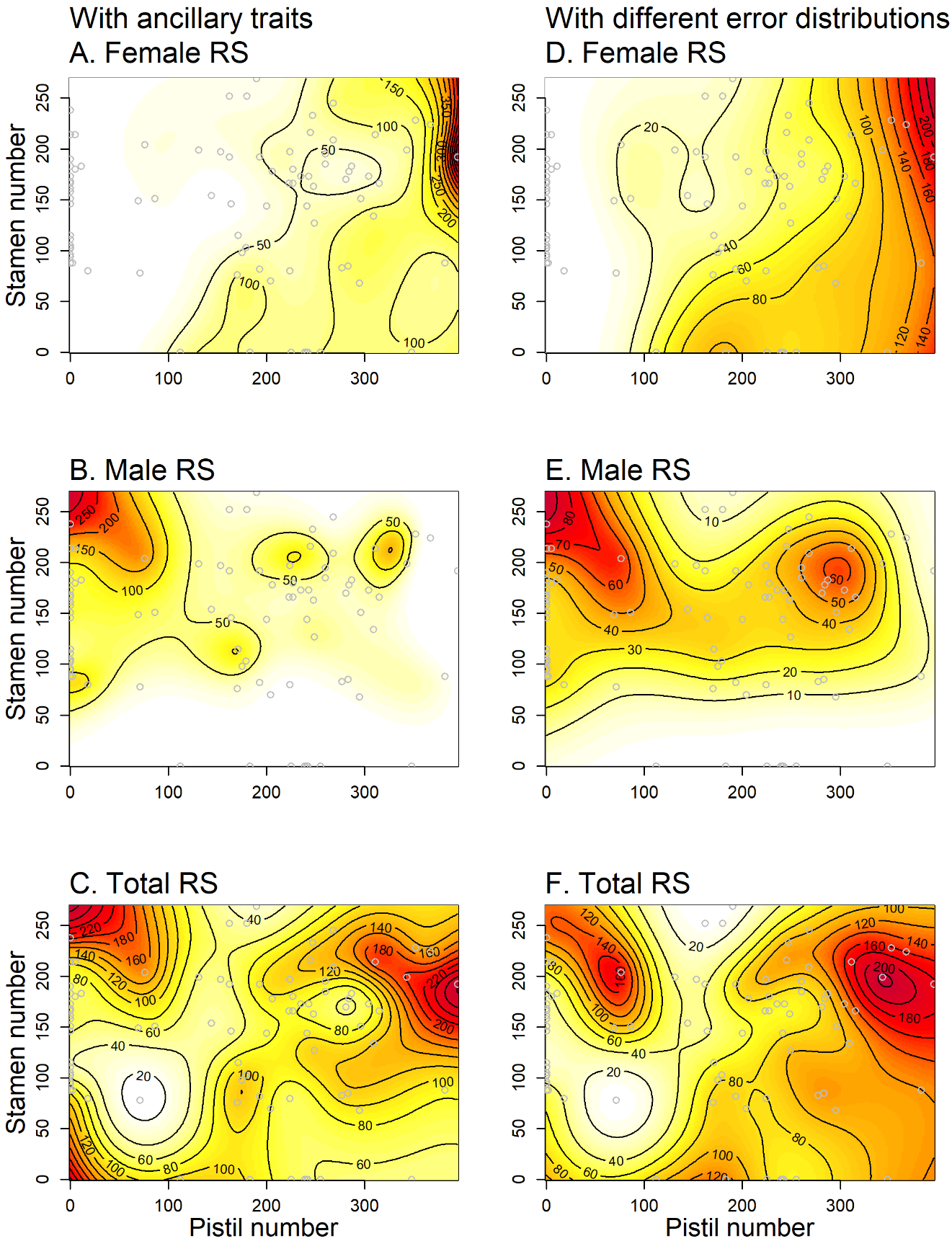

**Figure S6.** Representations of the fitness landscape for female, male, and total reproductive success (RS) as a function of pistil and stamen number in single-flowered individuals (*N* = 87; grey points), taking inbreeding depression into account in the supplementary analyses (related to STAR Methods). In the supplementary models, ancillary floral traits were included as covariates (**A**-**C**), or different error distributions were used (**D**-**F**). The colour gradient from white to red represents low to high predicted reproductive success as the number of seeds. In panels **A**-**C**, values are those predicted by generalized additive models (gam) using floral traits as covariates with a Poisson error distribution as the models presented in the main text. The floral traits, i.e., stalk height, petal length, and flowering date, were each fitted with a smoothing term of thin plate splines (with a basis dimension *k* = 3). In panels **D**-**F**, values are those predicted by generalized additive models (gam) using a negative binomial (for female and male RS) and zero-inflated Poisson (for total RS) distribution, with stamen and pistil number as two explanatory variables.

**Table S1.** Summary table of the shapes of fitness gain curves by nonlinear least square models under the scenario of an inbreeding depression of zero and 0.95. *P* values for the constant, *a*, indicate the statistical difference from zero, whereas *P* values for the exponent, *b*, indicate the statistical difference from one (linearity).

| **Inbreeding depression scenario** | **Sex function** | **Parameter** | **Estimate** | **Standard error** | ***t*-value** | ***P*** |
| --- | --- | --- | --- | --- | --- | --- |
| *δ* = 0 | Female | *a* | 0.01 | 0.01 | 1.11 | n.s. |
|  |  | *b* | 0.85 | 0.16 | 0.94 | n.s. |
|  | Male | *a* | 0.08 | 0.11 | 0.72 | n.s. |
|  |  | *b* | 0.52 | 0.27 | 1.77 | . |
| *δ* = 0.95 | Female | *a* | 3.7e-5 | 6.4e-5 | 0.59 | n.s. |
|  |  | *b* | 1.91 | 0.3 | 3.05 | ** |
|  | Male | *a* | 0.1 | 0.2 | 0.51 | n.s. |
|  |  | *b* | 0.47 | 0.38 | 1.38 | n.s. |

Notes: n.s. *P* > 0.1, . *P* < 0.1, ∗ *P* < 0.05, ∗∗ *P* < 0.01, ∗∗∗ *P* < 0.001

**Table S2.** Linear (β_i_), quadratic (γ_ii_), and correlational (γ_ij_) selection gradients on female and male allocation via female, male, and total reproductive success (RS) under the scenario of an inbreeding depression of zero and 0.95. Pistil and stamen number of each flower were standardized. *P* values for the estimates were calculated using *t*-tests in six multivariate regression models.

Notes: n.s. *P* > 0.1, . *P* < 0.1, ∗ *P* < 0.05, ∗∗ *P* < 0.01, ∗∗∗ *P* < 0.001

|  |  | **Female RS** |  | **Male RS** |  | **Total RS** | |  |
| --- | --- | --- | --- | --- | --- | --- | --- | --- |
| **Inbreeding depression scenario** | **Selection gradient** | **Estimate (SE)** | ***P*** | **Estimate (SE)** | ***P*** | **Estimate (SE)** | ***P*** | |
| *δ* = 0 | β (pistil number) | 0.72  (0.06) | *** | 0.19  (0.09) | * | 0.47  (0.06) | | *** |
|  | β (stamen number) | -0.031 (0.07) | n.s. | 0.24  (0.10) | * | 0.097  (0.07) | | n.s. |
|  | γ (pistil number) | -0.066 (0.07) | n.s. | -0.55  (0.10) | ** | -0.30  (0.07) | | * |
|  | γ (stamen number) | -0.029 (0.05) | n.s. | -0.43  (0.08) | ** | -0.22  (0.05) | | * |
|  | γ (pistil number, stamen number) | 0.19  (0.07) | n.s. | -0.084  (0.11) | n.s. | 0.059  (0.07) | | n.s. |
| *δ* = 0.95 | β (pistil number) | 0.82  (0.09) | *** | -0.23  (0.1) | ** | 0.41  (0.08) | | *** |
|  | β (stamen number) | -0.17  (0.1) | . | 0.36  (0.05) | *** | 0.03  (0.09) | | n.s. |
|  | γ (pistil number) | 0.7  (0.2) | ** | -39.7  (3.17) | *** | 0.4  (0.19) | | * |
|  | γ (stamen number) | 0.31  (0.15) | * | -8.13  (0.85) | *** | 0.05  (0.14) | | n.s. |
|  | γ (pistil number, stamen number) | 0.25  (0.21) | n.s. | -0.16  (0.42) | n.s. | 0.055  (0.19) | | n.s. |

**Table S3.** Estimated standardized linear (β_i_), quadratic (γ_ii_), and correlational (γ_ij_) selection gradients for pistil (p) and stamen (st) number via female, male, and total reproductive success (RS) for *Pulsatilla alpina* at the floral level from generalized additive models (*gam*). Standard errors and *P* values were obtained from bootstrapping procedures (see materials and methods for details).

Notes: n.s. *P* > 0.05, ∗ *P* < 0.05, ∗∗ *P* < 0.01, ∗∗∗ *P* < 0.001

|  |  | **Female RS** |  | **Male RS** |  | **Total RS** |  |
| --- | --- | --- | --- | --- | --- | --- | --- |
| **Inbreeding depression scenario** | **Selection gradient** | **Estimate (SE)** | ***P*** | **Estimate (SE)** | ***P*** | **Estimate (SE)** | ***P*** |
| *δ* = 0 | β _p_ | 0.57  (0.03) | *** | 0.0098  (0.05) | n.s. | 0.34  (0.02) | *** |
|  | β _st_ | -0.10  (0.02) | *** | 0.27  (0.03) | *** | 0.010  (0.02) | n.s. |
|  | γ _p_ | -5.78  (0.92) | *** | -15.17  (1.40) | *** | -4.41  (0.51) | *** |
|  | γ _st_ | -1.9  (0.26) | *** | -2.04  (0.32) | *** | -1.21  (0.13) | *** |
|  | γ _p, st_ | 0.15  (0.13) | * | -0.74  (0.2) | n.s. | -0.13  (0.07) | n.s. |
| *δ* = 0.95 | β _p_ | 0.8  (0.04) | *** | -0.23  (0.1) | *** | 0.35  (0.04) | *** |
|  | β _st_ | -0.11  (0.03) | *** | 0.36  (0.05) | *** | 0.04  (0.03) | n.s. |
|  | γ _p_ | -2.76  (0.96) | *** | -39.69  (3.05) | *** | -6.95  (0.82) | *** |
|  | γ _st_ | -1.66  (0.27) | *** | -8.13  (0.84) | *** | -2.07  (0.23) | *** |
|  | γ _p, st_ | -0.25  (0.17) | n.s. | -0.16  (0.42) | n.s. | -0.33  (0.1) | *** |
